## Supplementary Note 1, Supplementary Figures 1-19, Supplementary Tables 1-4 for "Rational engineering of allosteric protein switches by *in silico* prediction of domain insertion sites"

**Contents:**

Supplementary Note 1
Supplementary Figures 1-19
Supplementary Tables 1-4

**Supplementary Note 1 | Details on model training and optimization.**

Having built a large domain insertion dataset, we aimed to employ it for machine learning to predict sites in proteins suitable for domain insertion and thus for protein engineering. To this end, we removed the insert domain sequence portion from each protein in our data set. This resulted in semi-synthetic sequences of proteins lacking their naturally occurring inserts (parent_only). Importantly, for all parent_only proteins, we have information about the true domain insertion site. This site and its two adjacent positions were considered as positive labels for model training, since exact domain boundaries are difficult to define and previous experimental work has shown that domain insertions are often tolerated at several consecutive sequence positions[1,2]. All other sequence positions were assumed not to tolerate domain fusion and were therefore assigned negative labels. Importantly, this assumption is rather naive, as some of these positions may, in fact, tolerate domain fusion (see below). That said, due to the lack of further information on these sites and in light of experimental evidence that most positions in proteins do not tolerate domain fusion[1,2], this label assignment is a reasonable basis for model training (Fig. 2a).

Subsequently, the dataset was randomly split into train validation and test sets (ratio 70:20:10; referred to as “random” split). Since simple one-hot encoding of sequences was, as expected, not suitable for model training (Fig. 2b and Supplementary Fig. 3a), we computed information-rich multidimensional representations (embeddings) of the parent_only protein sequences with ESM-2[3]. Based on these embeddings, we then trained multi-layer perceptron (MLP) decoders to identify the true domain insertion sites in the parent_only sequences. The resulting model accurately inferred true insertion sites with a median probability score of 0.9, while reliably assigning low scores to negatively labeled positions (median score = 4.32e-06; Fig. 2b and Supplementary Fig. 3b). We further investigated whether scaling the decoder to more complex architectures would yield a significant performance improvement. Indeed, replacing the MLP with BERT architectures of either 36.7 or 146.0 million parameters improved discrimination between positive and negative labels, mainly due to increased prediction scores for the positive positions[4] (Supplementary Fig. 3b).

The near-perfect performance on the validation set raised the question of whether our model had simply memorized information about suitable insertion sites between different members of the same domain superfamily present in the training and test sets, regardless of the strict sequence similarity cutoff we had used. To enforce generalization beyond the training data, we first split the data set by Interpro/CATH superfamily terms (referred to as “Interpro” split), meaning that members of the same superfamily occur in only one of the three subsets. To create an even more stringent dataset split, we randomly selected only one example sequence for each parent domain superfamily, resulting in a much smaller dataset of only 232 highly diverse sequences (referred to as “single” split). We trained our MLP decoder again on these new datasets and compared all three training regimes on an intersected test set (Fig. 2c, Supplementary Fig. 3b). While we observed a greater overlap between the predicted probability distributions, both new models were generally still able to discriminate between insertion tolerant and non-tolerant sites (Fig. 2c).

To further optimize our approach, we considered two unique features of our dataset. First, the data is by definition highly unbalanced, with only three consecutive positive sites per parent_only sequence, which biases any model toward global prediction of low values (i.e. a trivial model that predicts only zeros has a high overall success rate). Second, we only have reliable information about the positive labels, whereas we do not know whether domain insertions would in fact be tolerated at some sites that are assumed to be negative.

To account for these factors, we introduced a masking strategy during loss computation that considers only the positive labels and a single, randomly selected, negatively labeled site of each parent_only sequence in each training epoch. This approach significantly increases the weight of the positive labels. Importantly, as the negative site is randomly selected in each epoch, the model still captures information about the entire protein sequence as training progresses. We assumed that this approach would facilitate the model to transfer the patterns learned on the true positive sites to the unknown protein regions, since the model would be less penalized in this regard during training. When we tested this strategy on all three dataset splits (random, Interpro, single), we observed successful discrimination between positive and negative labels in all cases (Fig. 2d). Importantly, models based on both the Interpro and single superfamily data splits, robustly predicted high scores for the true positives (median prediction of 0.46 (Interpro) and 0.87 (single) compared to 0.002 and 0.3 for negative labels, respectively; compare Fig. 2c and 2d). As intended, the model predicted approximately 10% of sequence positions as positives in regions of the parent_only protein, now including sites for which we had no information on domain insertion tolerance. The frequency of positives is also in line with our expectation derived from previous experimental observations[1,2]. We selected our model trained on the most restrictive, single sequence data split for further investigation and termed it ProDomino.

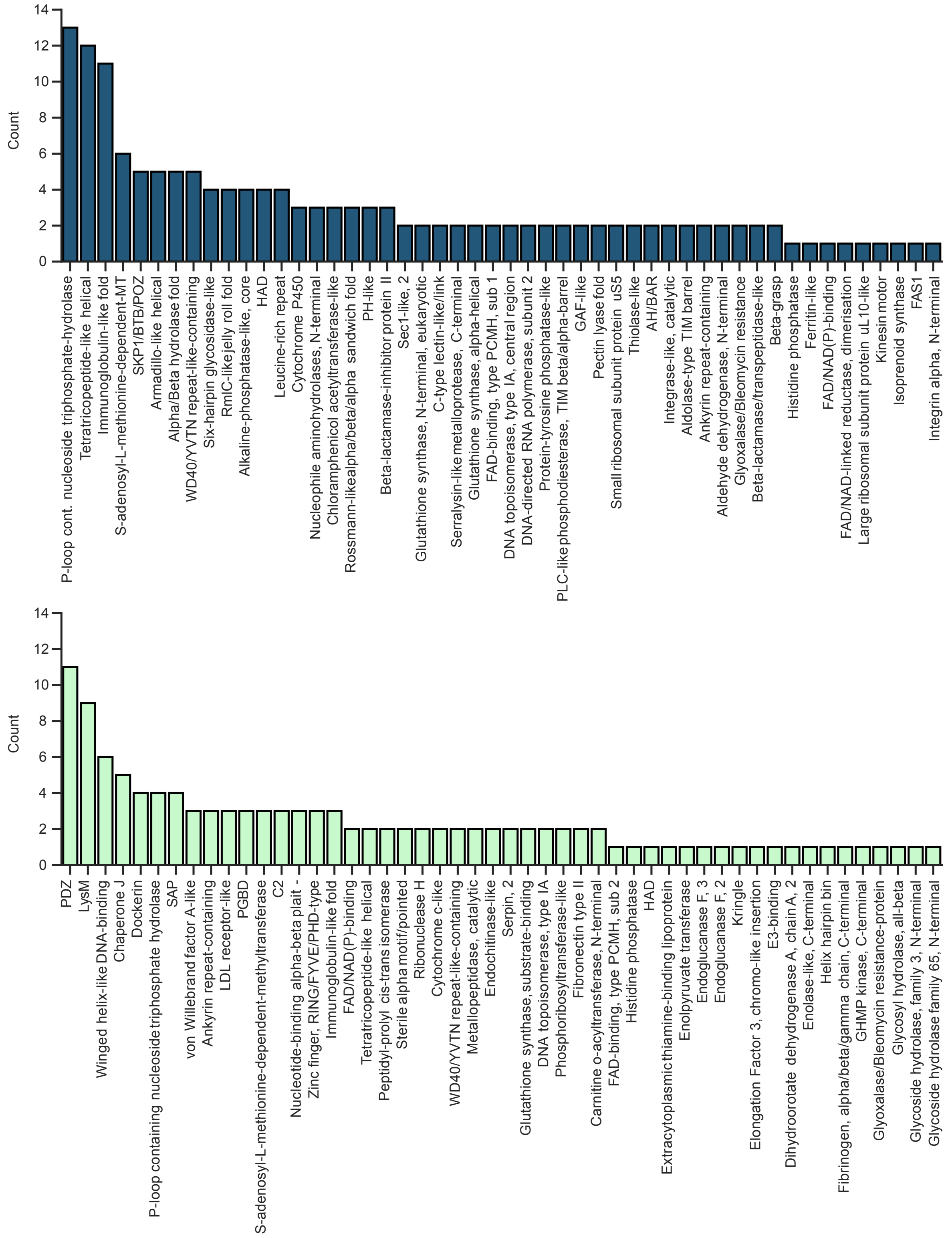


**Supplementary Fig. 1 | Parent and insert domains show different degrees of promiscuity with respect to their fusion partners.** The number of unique domain superfamily combinations is shown for a given parent (top panel) or insert domain (bottom panel) for top 50 domain superfamilies. The data correspond to Fig. 1c.

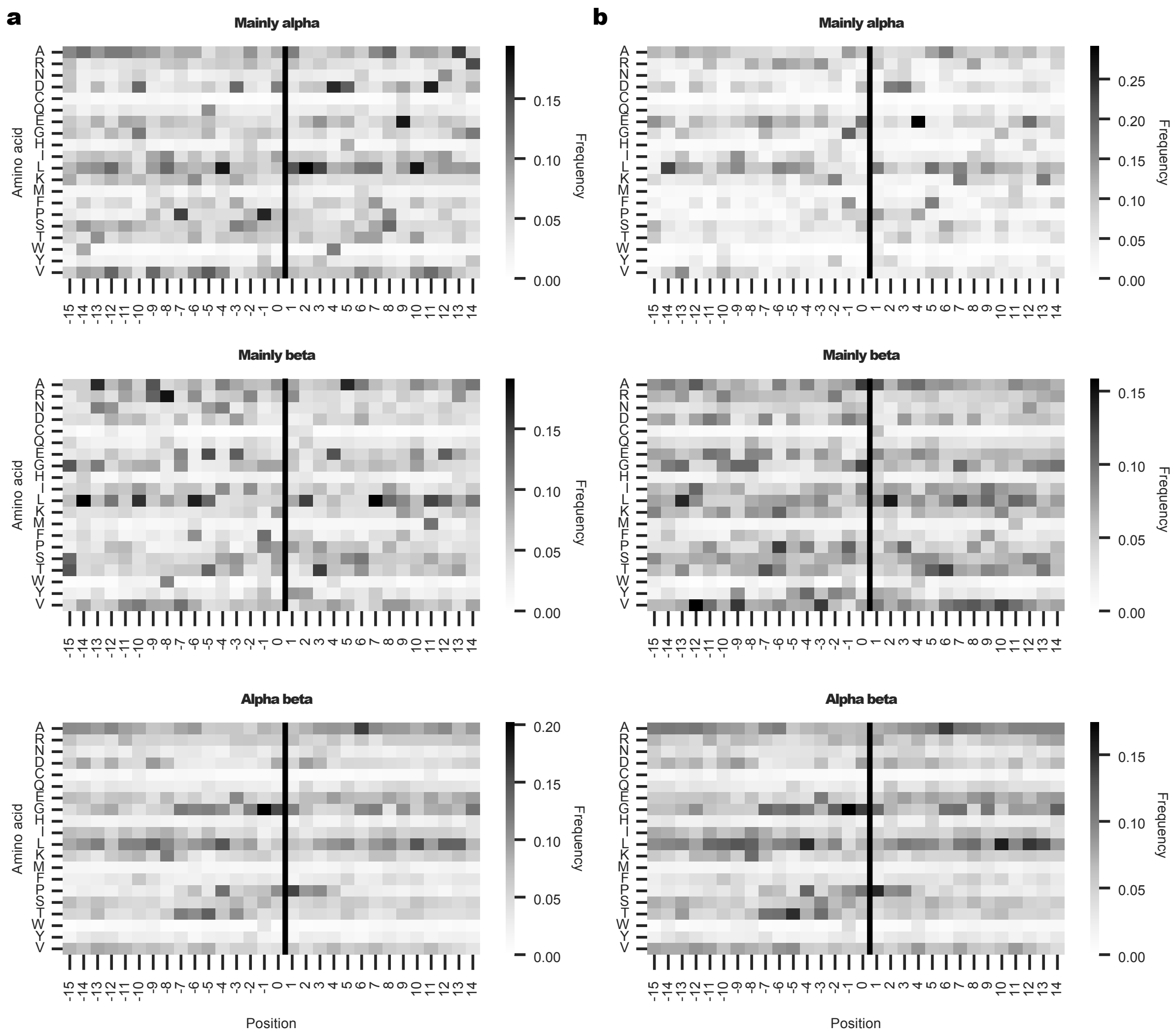


**Supplementary Fig. 2 | Natural domain insertion sites do not show enrichment for specific sequence motifs. a**, **b** Heatmaps of amino acid frequencies surrounding domain insertion sites are shown for different protein groups within the dataset. Heatmaps were generated for all parent domains belonging to the indicated CATH class (mainly alpha, mainly beta or alpha-beta) (**a**) and for all inserts within these classes (**b**). Black lines indicate domain insertion sites.

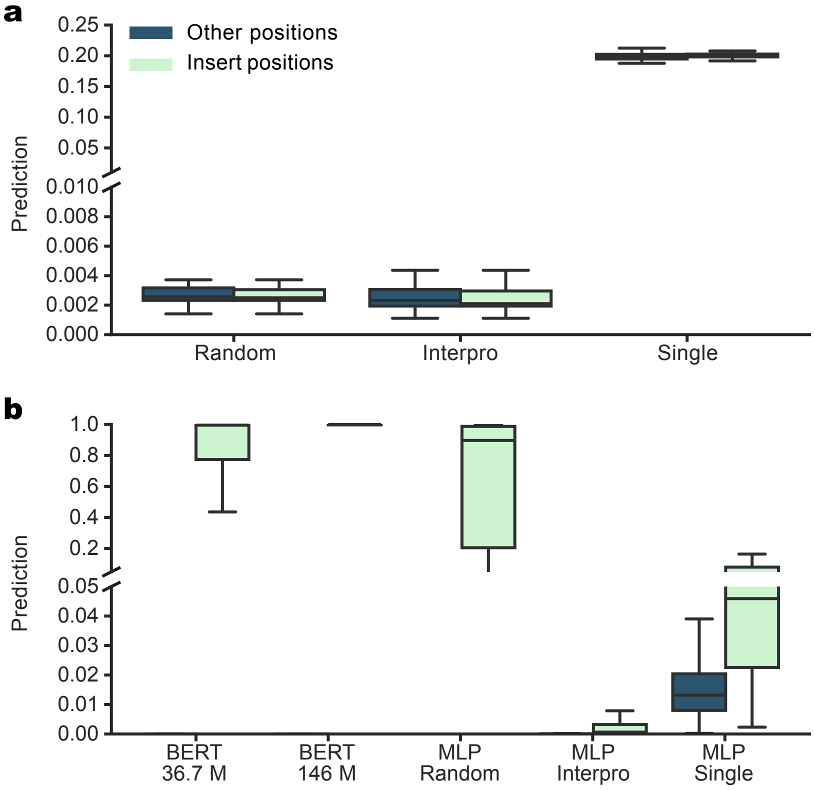


**Supplementary Fig. 3 | Optimization of ProDomino models. a,** Box plots show the test set performance of additional baseline MLP models trained on one-hot encodings (corresponds to Fig. 2c). **b**, The performance of BERT-like classifiers on the test set is shown. Box plots of the corresponding MLP classifiers from Fig. 2c are shown for comparison. **a**,**b** Boxes represent the interquartile range (IQR) and the median is represented by a horizontal line. Whiskers extend to the 1.5-fold IQR or to the value of the smallest or largest prediction value.

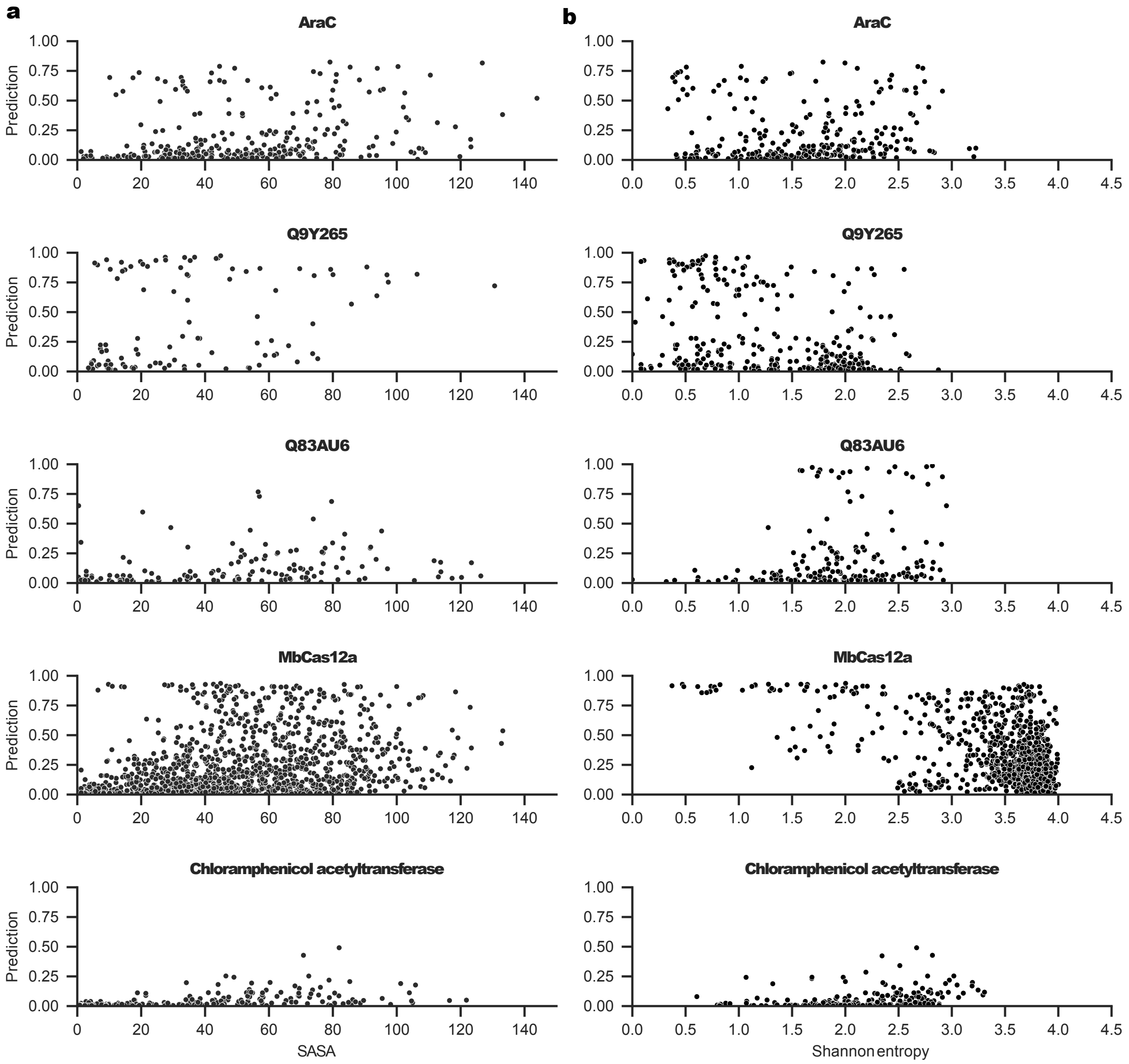


**Supplementary Fig. 4 | ProDomino predictions do not correlate with SASA or sequence conservation. a**, Scatterplots of the relation between ProDomino predictions and SASA for sample proteins with available structures. **b**, Scatterplots of the relation between ProDomino predictions and sequence conservation, represented by Shannon entropy, for selected proteins.

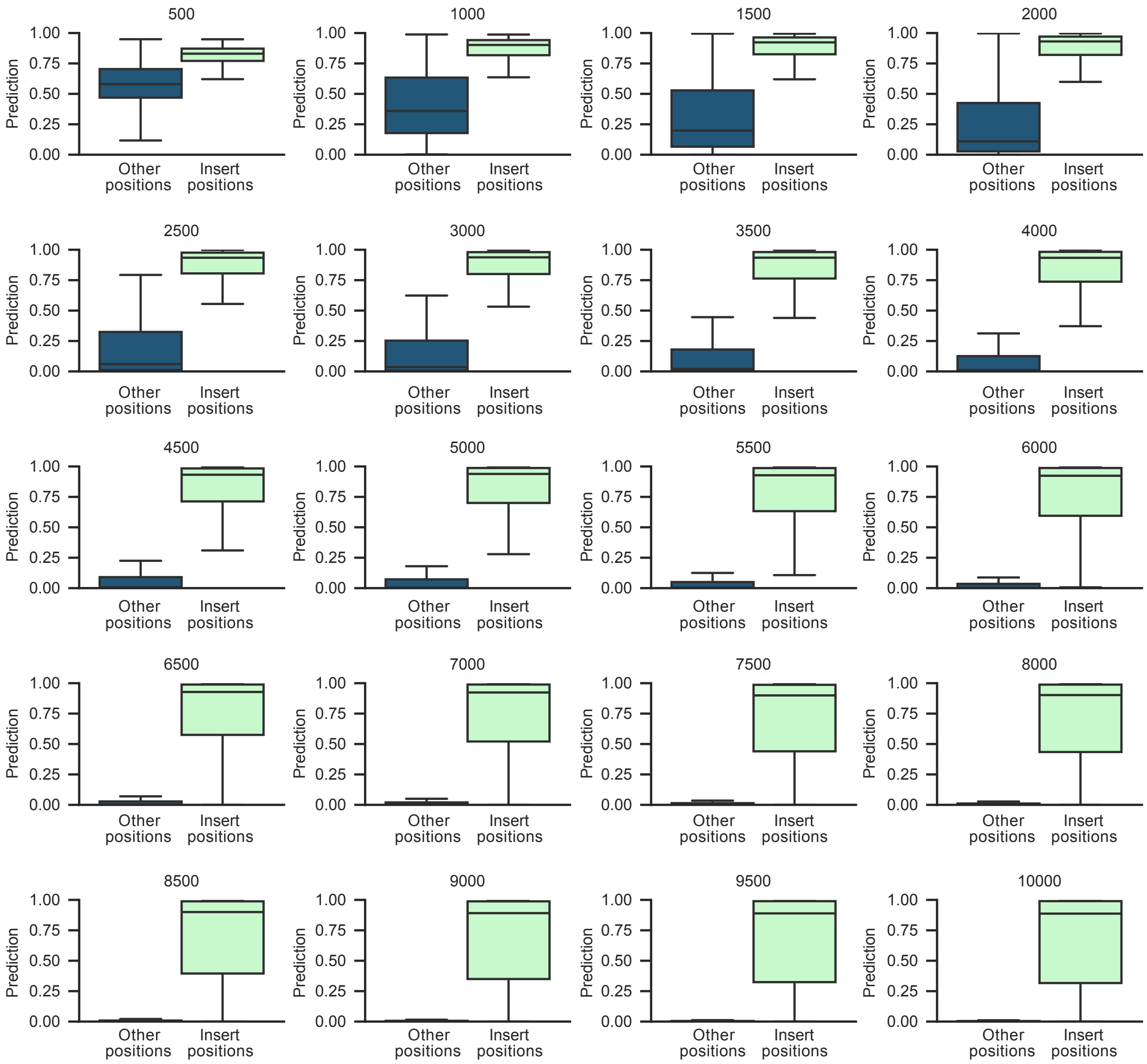


**Supplementary Fig. 5 | The number of training steps affects model sensitivity.** Model prediction scores for true insertion sites and other (unknown) positions are shown as box plots. Numbers above each plot indicate model training duration in steps. Boxes represent the interquartile range (IQR) and the median is represented by a horizontal line. Whiskers extend to the 1.5-fold IQR or to the value of the smallest or largest predicted value.

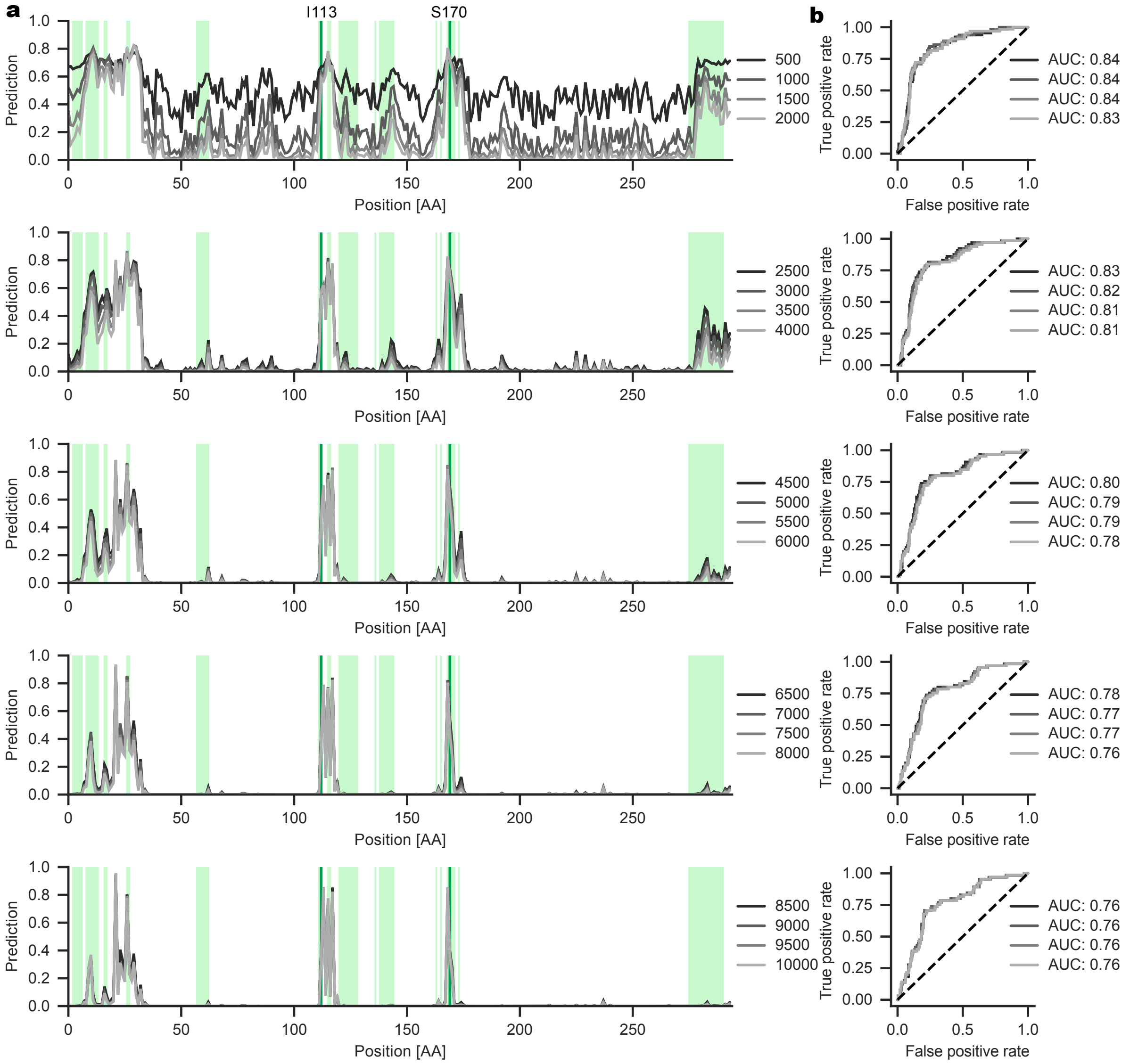


**Supplementary Fig. 6 | ProDomino correctly identifies insertion-tolerant regions in AraC. a**, The insertion score for the bacterial transcription factor AraC is shown for each amino acid position. The scores of models trained for different numbers of steps are shown in different shades of gray. Five individual subplots are presented for clarity. Green regions indicate experimentally validated insertion tolerant sites. The two sites previously used to engineer light-regulated AraC variants, I113 and S170, are indicated in dark green. **b**, ROC curves based on the predictions in **a** are shown. The area under the curve (AUC) is given for each model variant.

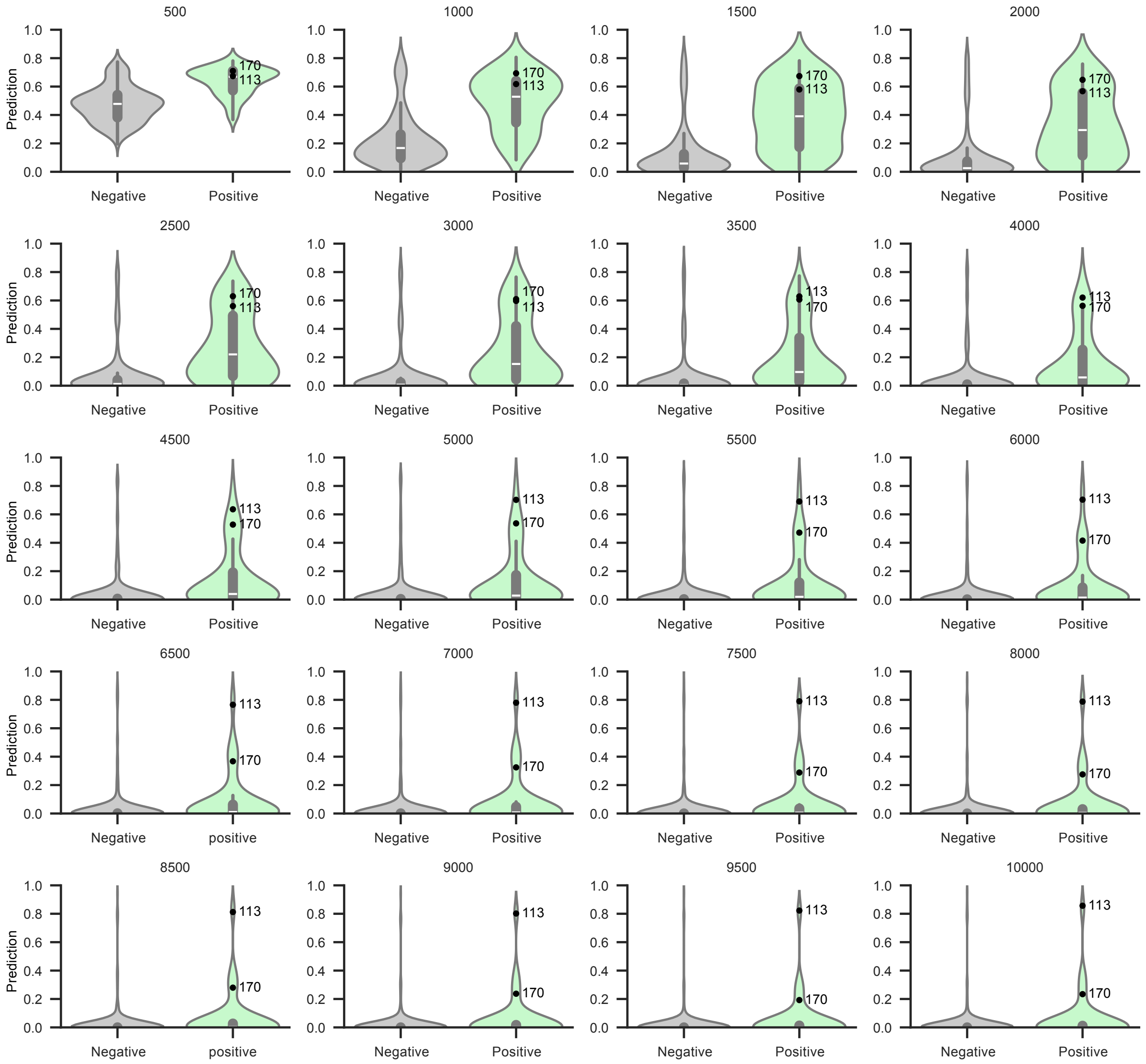


**Supplementary Fig. 7 | ProDomino identifies allosteric sites previously used to engineer switchable AraC variants.** Violin plots showing the distribution of predictions for insertion tolerant sites (green) and positions not amenable to domain insertion (gray). The scores for the insertion sites I113 and S170, which are known to give rise to light-regulated AraC variants when a LOV2 photosensory domain is introduced, are indicated. Boxes represent the interquartile range (IQR) and the median is represented by a white horizontal line. Whiskers extend to the 1.5-fold IQR or to the value of the lowest or highest predicted value.

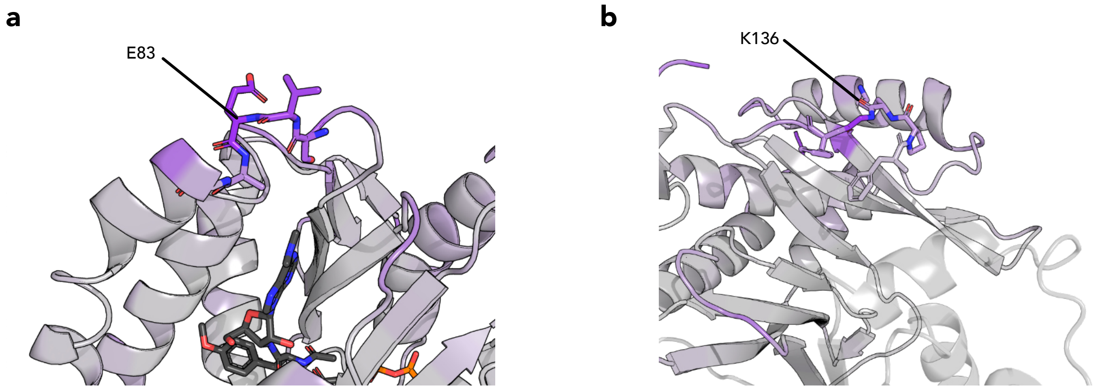


**Supplementary Fig. 8 | Detailed view of the insertion sites of the lead PAC and CAT candidates. a**, **b**, ProDomino inferred insertion scores are mapped onto the crystal structures of PAC (**a**) and CAT (**b**). The insertion sites of the lead candidates are indicated and shown in the side chain representation. PDB ID s: 7K0A, 1PD5.

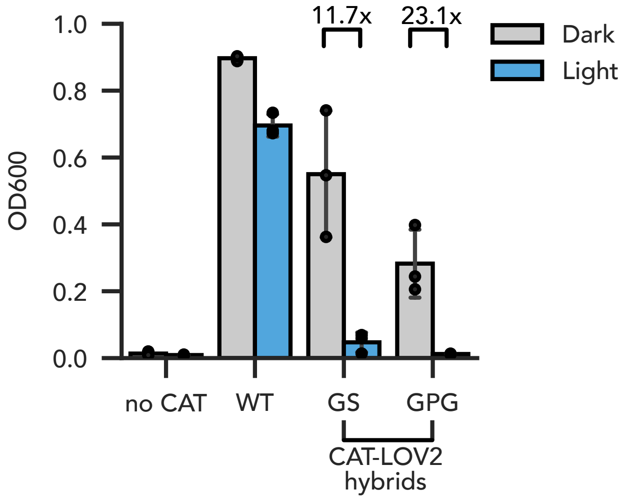


**Supplementary Fig. 9 | Light-controlled *E. coli* cell growth.** Bacteria were transformed with plasmids expressing the indicated CAT variant (insertion behind K136) or an empty control plasmid. Liquid cultures were grown in the presence of 50 µg/mL chloramphenicol for 7 hours and cell density was assessed by measuring OD at 600 nm. Bars indicate means, error bars the standard deviation, and black dots individual data points from n=3 independent experiments.

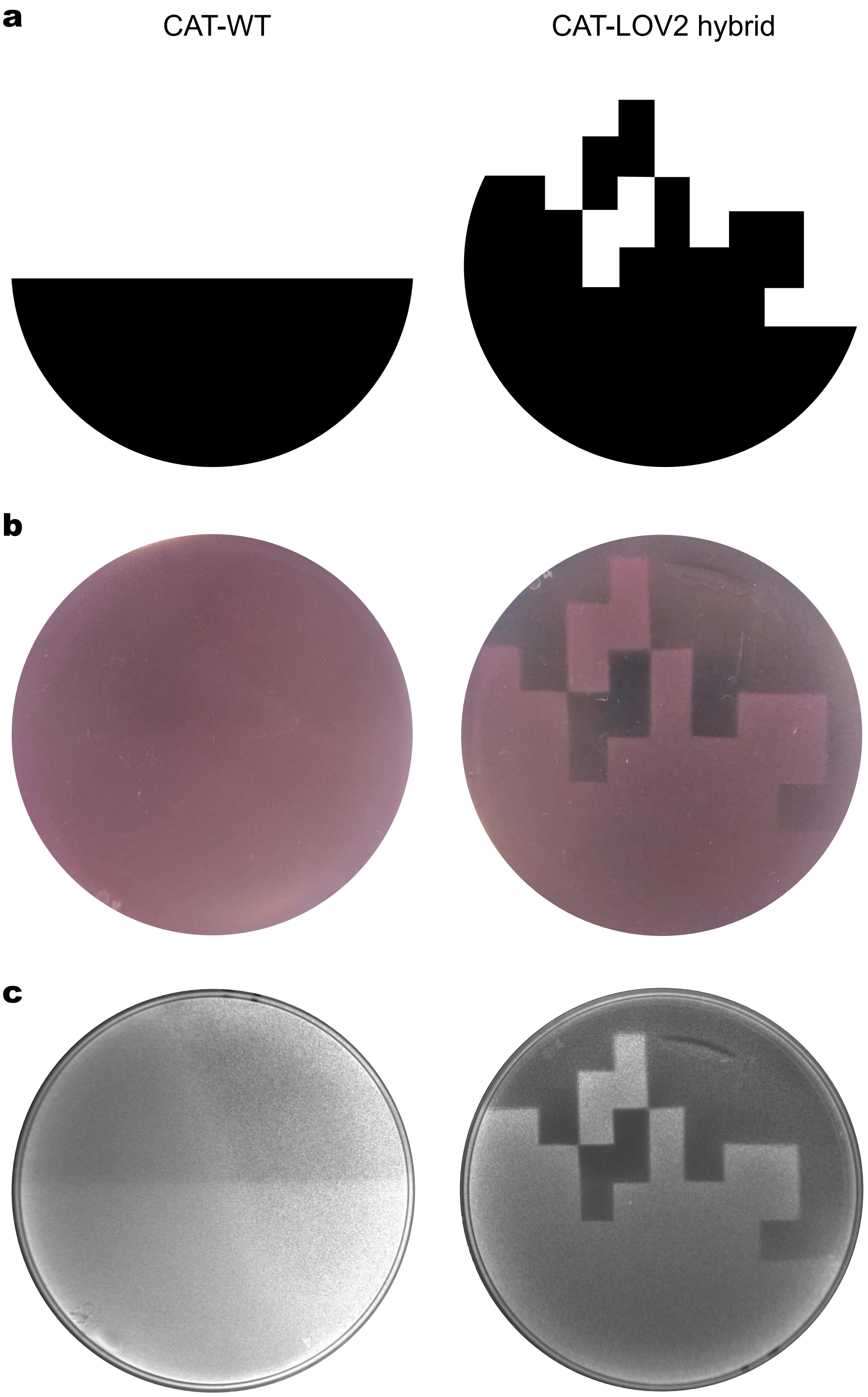


**Supplementary Fig. 10 | Spatial control of *E. coli* growth. a-c,** *E. coli* expressing CAT-K136-LOV or a constitutively active wild-type CAT and mRFP (for visualization) were plated in top agar supplemented with 25 µg/mL chloramphenicol. During incubation at 37°C, the plates were illuminated through the depicted photomask (**a**) and mRFP was imaged under white light (**b**) or UV light (**c**). The lower right image corresponds to Fig. 3h.

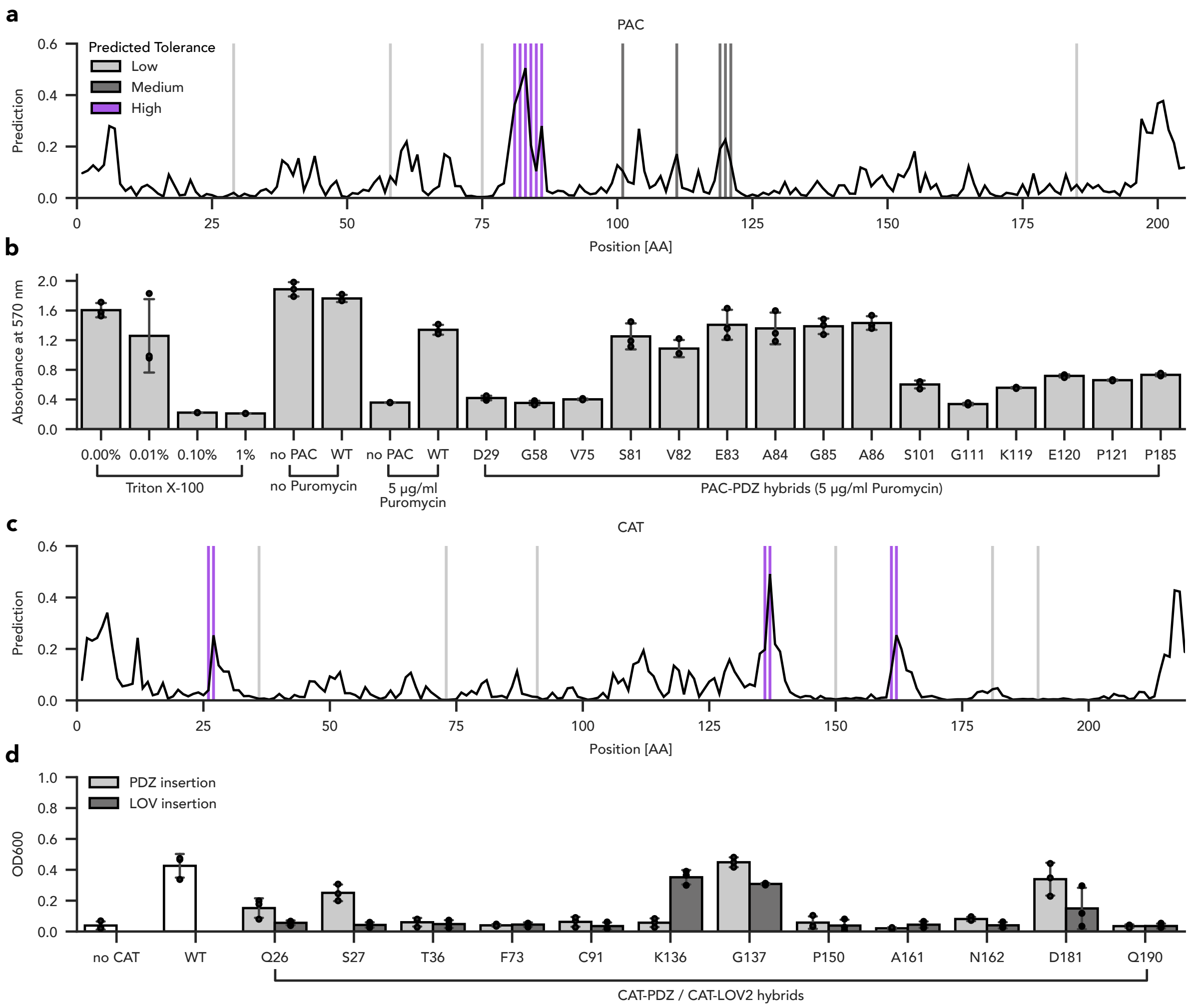


**Supplementary Fig. 11 | Domain insertion screening of PAC and CAT confirms ProDomino predictions. a, c,** ProDomino inferred insertion scores are mapped onto the primary sequence of PAC (**a**) and CAT (**c**). Insertion sites selected for experimental testing are marked by vertical lines and color coded as indicated. **b**, Assessment of insertion tolerance in PAC. HEK293T cells were transfected with vectors encoding the respective PAC variants carrying PDZ insertions after the indicated residue or a negative control expressing enhanced green fluorescent protein (eGFP). Cells were treated with 5 µg/mL puromycin and incubated for 48 hours before cell viability was assessed by MTT assay. Cells treated with different concentrations of toxic Triton X-100 served as controls for the assay itself. Bars indicate means, error bars the standard deviation, and black dots individual data points from n=3 independent experiments. **d**, Assessment of the CAT insertion permissibility. Bacteria were transformed with plasmids expressing the indicated CAT variant or an empty control plasmid. Liquid cultures were grown in the presence of 25 µg/mL chloramphenicol for 7 hours and cell density was assessed by measuring OD at 600 nm. Light gray bars represent PDZ insertions behind the indicated residue and dark gray bars correspond to LOV2 insertions at the same position. Bars indicate means, error bars the standard deviation, and black dots individual data points from n=3 independent experiments.

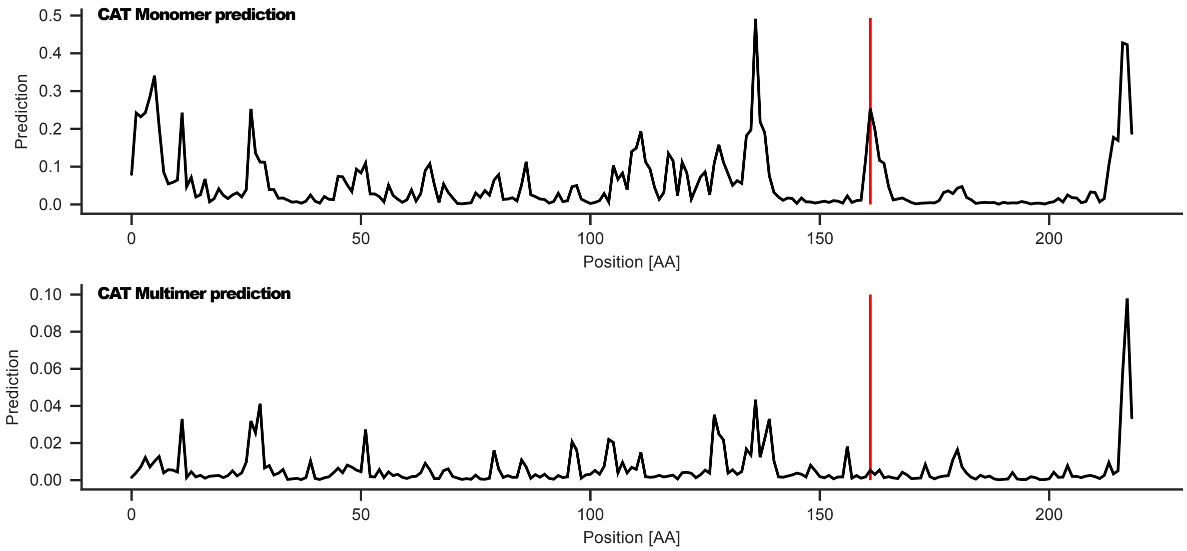


**Supplementary Fig. 12 | Insertion score prediction for an artificially concatenated, trimeric CAT correctly identifies a false positive site.** ProDomino predictions for CAT are shown based on the wild-type sequence (top panel) or a concatenated 3xCAT sequence (bottom panel) as input. The red line marks the “false positive” site, which we observed not to tolerate PDZ insertion (see Supplementary Fig. 11d).

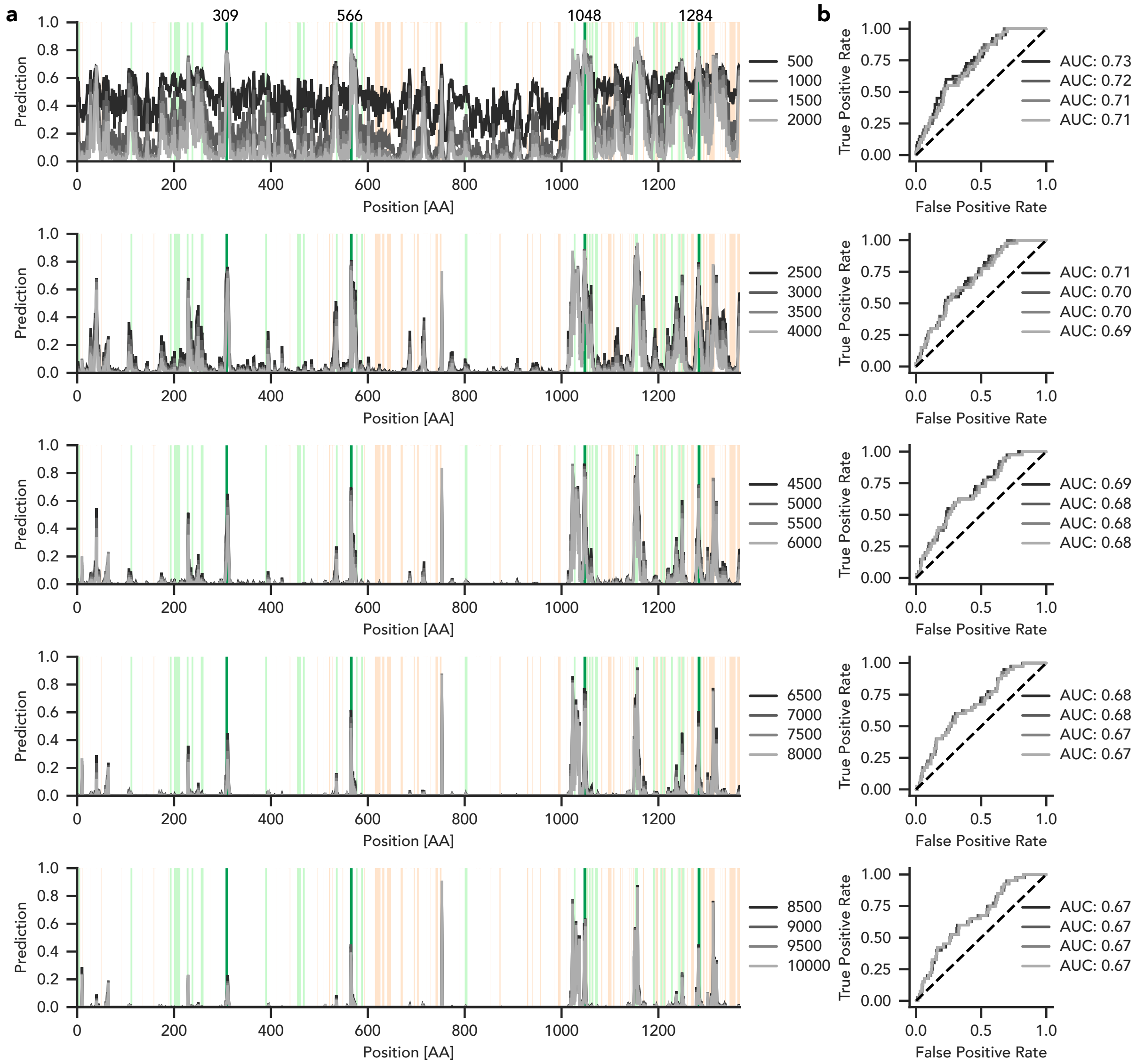


**Supplementary Fig. 13 | ProDomino predictions correlate with insertion tolerant regions in Cas9. a**, The insertion score for Cas9 is shown for each amino acid position. The scores of models trained for different numbers of steps are shown in different shades of gray. Regions in light green indicate insertion permissive sites, orange marks sites that were not amenable to domain fusion, as reported by Oakes et al.[5]. Positions that have been experimentally validated by us (Fig. 4c) are marked in dark green. **b**, ROC curves based on the predictions in **a** are shown. The area under the curve (AUC) is given for each model.

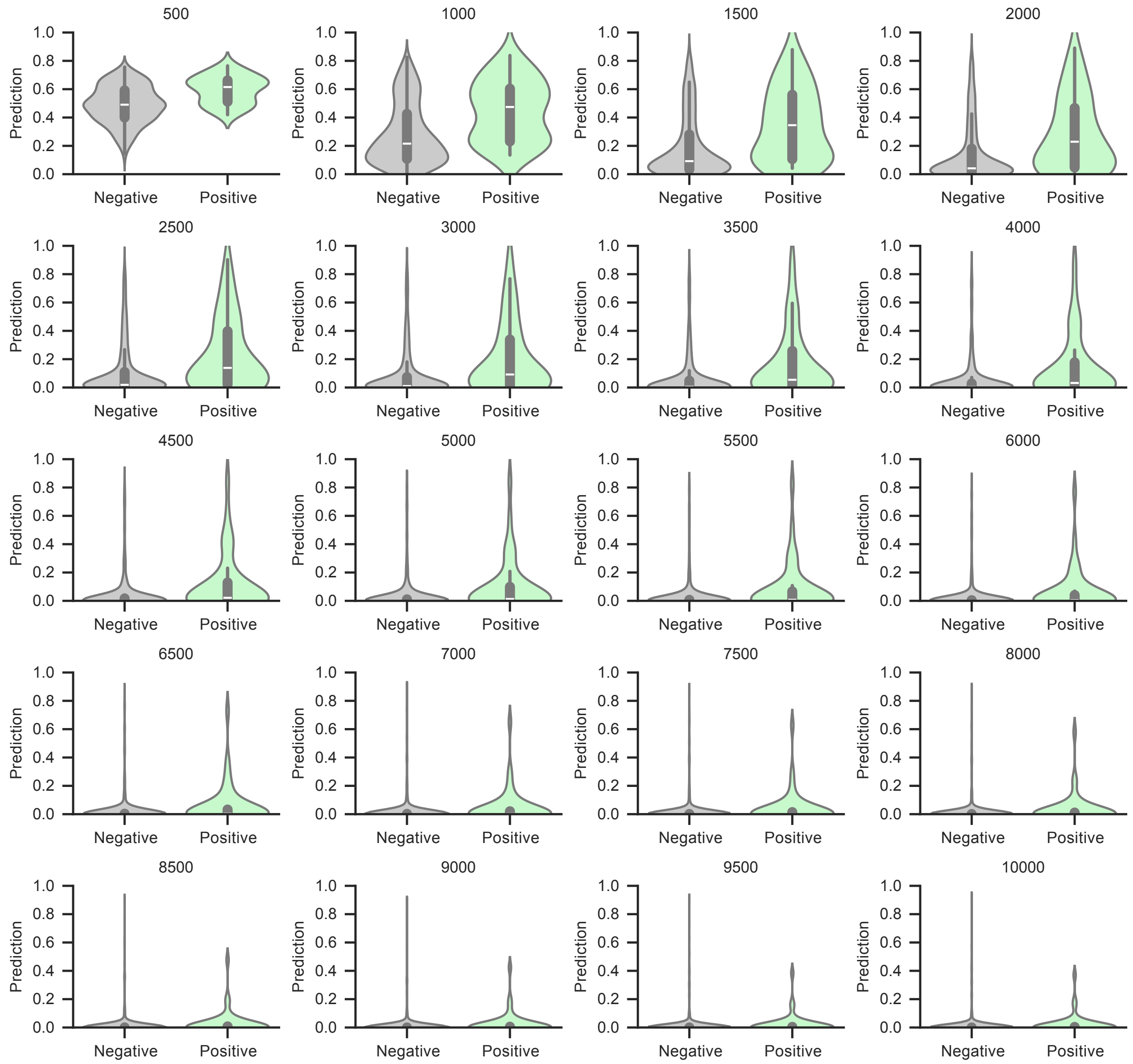


**Supplementary Fig. 14 | Comparison of prediction score distributions for insertion permissive or non-permissive sites in Cas9.** Violin plots showing the distribution of ProDomino predictions for insertion permissive sites (green) and non-permissive sites (gray). Positive and negative labels correspond to the data reported by Oakes et al.[5]. Boxes represent the interquartile range (IQR) and the median is represented by a white horizontal line. Whiskers extend to the 1.5-fold IQR or to the value of the smallest or largest prediction value.

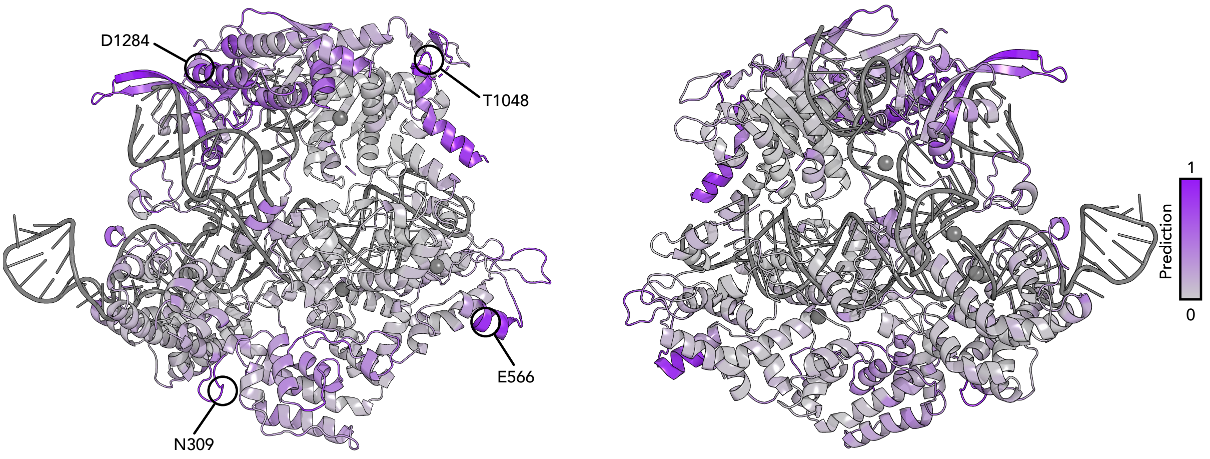


**Supplementary Fig. 15 | ProDomino prediction scores mapped onto the Cas9 structure.**
Insertion scores correspond to the 1,500-step model in Supplementary Fig. 13. PDB ID: 4UN3.

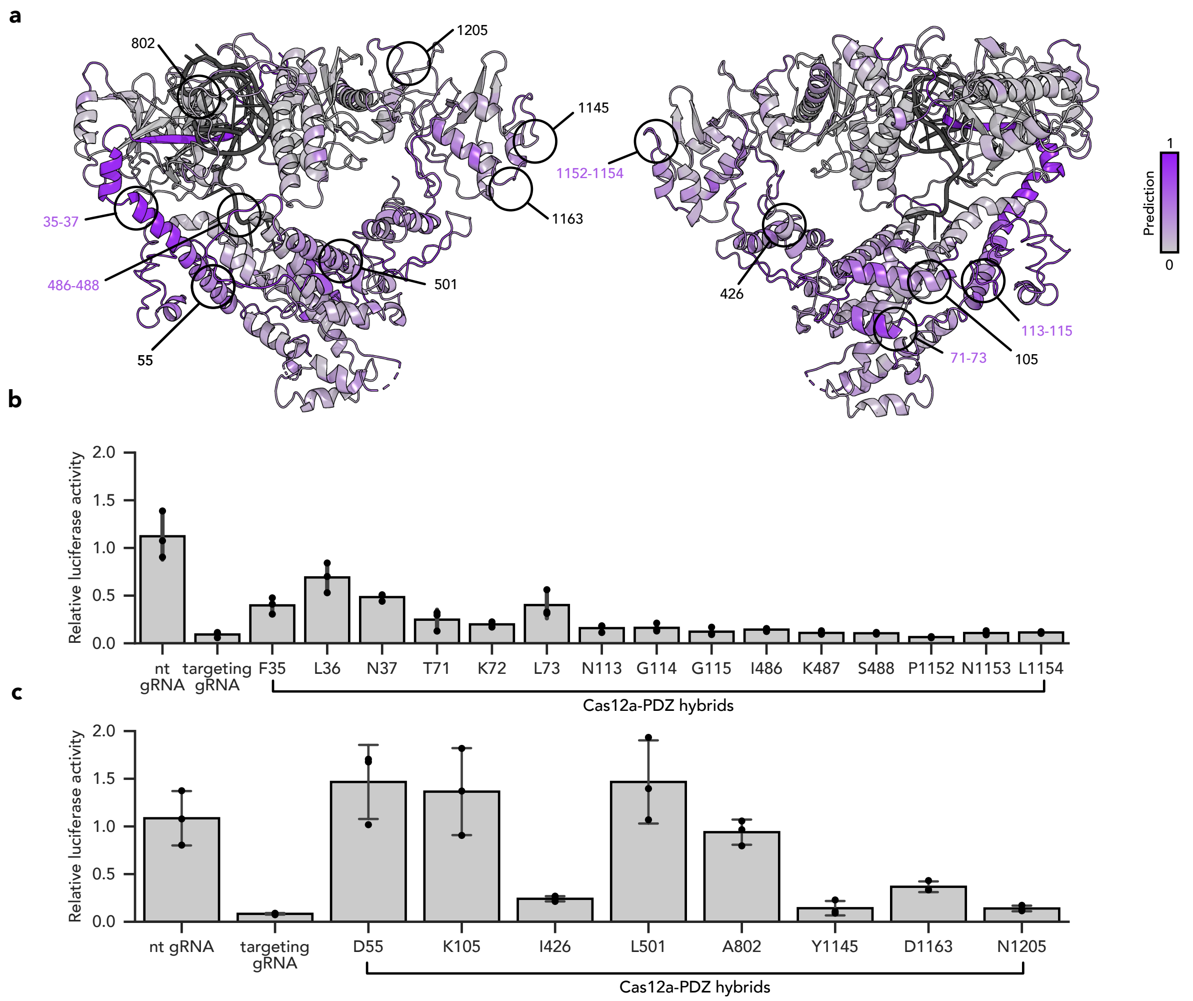


**Supplementary Fig. 16 | Experimental assessment of insertion tolerance in Cas12a. a**, Insertion scores are mapped onto a cryo-electron microscopy (cryo-EM) structure of *Mb*Cas12a. PDB ID: 6IV6. **b**, **c**, HEK293T cells were transfected with vectors encoding (i) the indicated Cas12a-PDZ insertion variant, (ii) a firefly luciferase targeting gRNA and (iii) a luciferase reporter. Samples were incubated for 48 hours, and luciferase activity was measured in a plate reader. The activity of insertion variants predicted to be active (**b**) or inactive (**c**) is shown. Bars indicate means, error bars the standard deviation, and black dots individual data points from n=3 independent experiments. nt, non-targeting gRNA

**a**

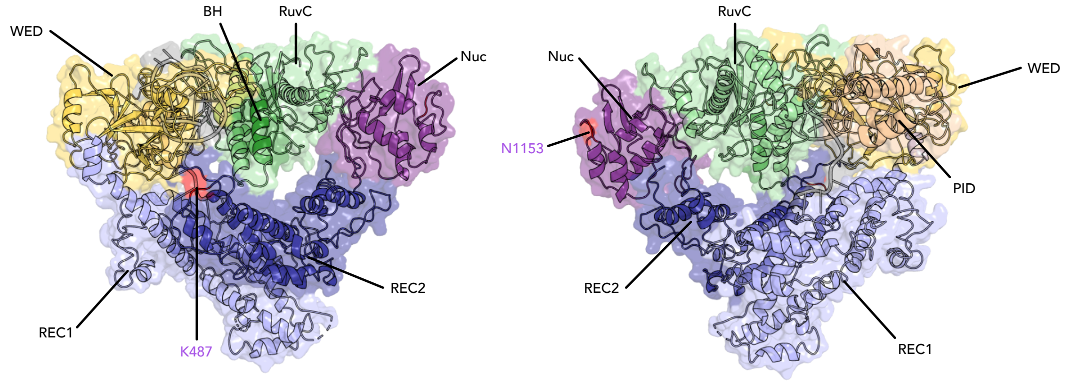


**Supplementary Fig. 17 | Positions of the two lead insertion sites within Cas12a.** The cryo-EM structure of Cas12a is shown in two orientations. The two allosteric insertion sites, K487 and N1153, are marked in red. The individual domains of Cas12a are indicated and color-coded. WED: wedge; BH: bridge helix; Nuc: nuclease; REC: recognition; PID: PAM interacting domain. PDB ID: 6IV6.

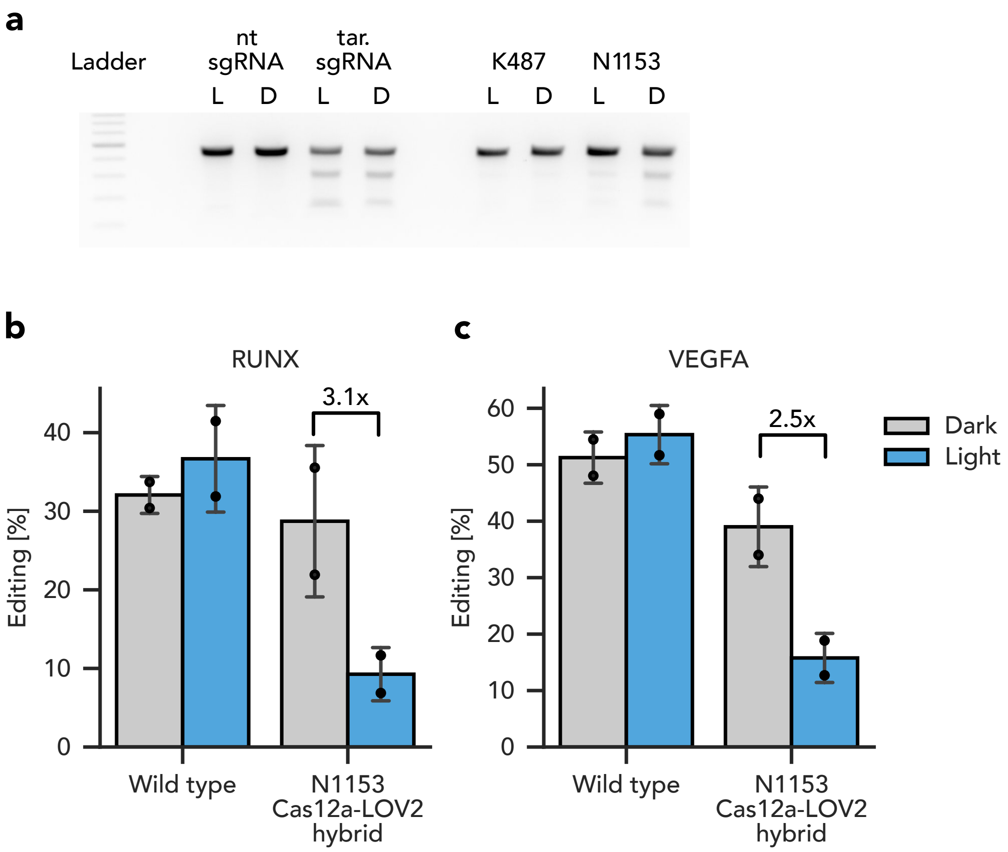


**Supplementary Fig. 18 | Light-dependent regulation of *Mb*Cas12a.**

**a-c,** HEK293T cells were transfected with vectors encoding (i) the indicated Cas12a variant and (ii) a gRNA targeting the endogenous *RUNX* (**a**, **b**) or *VEGFA* (**c**) locus. Samples were incubated under blue light illumination or in the dark for 72 h. InDel frequencies were assessed qualitatively via T7E1-assay (**a**) or quantitatively by NGS (**b**, **c**). A representative agarose gel image is shown in **a**. (**b,c**) Bars indicate means, error bars the standard deviation, and black dots individual data points from n=2 independent experiments.

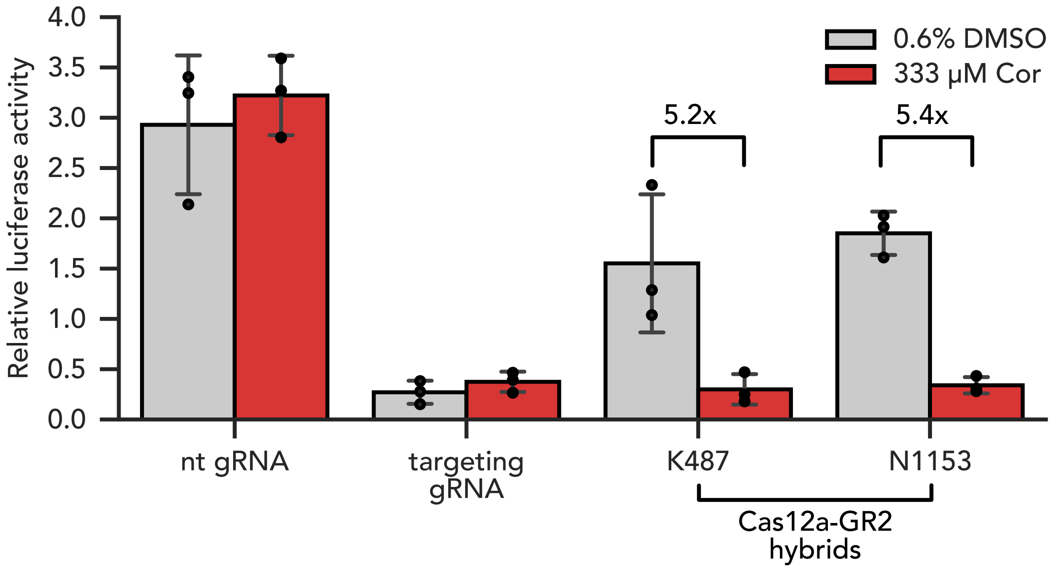


**Supplementary Fig. 19 | Tight, chemical regulation of Cas12a.** HEK293T cells were transfected with vectors encoding (i) the indicated Cas12a variant, (ii) a firefly luciferase gene targeting gRNA and (iii) a dual luciferase reporter. The Cas12a hybrids carried the GR2 receptor domain as insert at positions K487 or N1153. Samples were treated with cortisol or DMSO (solvent) as indicated. At 48 hours post-transfection, luciferase activity was measured in a plate reader. Bars indicate means, error bars the standard deviation, and black dots individual data points from n=3 independent experiments. Cor, cortisol.

**Supplementary Table 1 | List of constructs used in this study.** Insert domains are flanked by SG linkers on both sides, unless otherwise indicated. WT, wild type; CMV, cytomegalovirus; eGFP, enhanced green fluorescent protein.

| # | Name | Description, sequential order |
|---|---|---|
| 1 | BLA_CAT | β-lactamase expression cassette, chloramphenicol acetyltransferase expression cassette |
| 2 | BLA | β-lactamase expression cassette |
| 3 | BLA_CAT_F25_PDZ | Chloramphenicol acetyltransferase with a PDZ insertion behind F25 |
| 4 | BLA_CAT_Q26_PDZ | Chloramphenicol acetyltransferase with a PDZ insertion behind Q26 |
| 5 | BLA_CAT_S27_PDZ | Chloramphenicol acetyltransferase with a PDZ insertion behind S27 |
| 6 | BLA_CAT_T36_PDZ | Chloramphenicol acetyltransferase with a PDZ insertion behind T36 |
| 7 | BLA_CAT_F73_PDZ | Chloramphenicol acetyltransferase with a PDZ insertion behind F73 |
| 8 | BLA_CAT_C91_PDZ | Chloramphenicol acetyltransferase with a PDZ insertion behind C91 |
| 9 | BLA_CAT_P135_PDZ | Chloramphenicol acetyltransferase with a PDZ insertion behind P135 |
| 10 | BLA_CAT_K136_PDZ | Chloramphenicol acetyltransferase with a PDZ insertion behind K136 |
| 11 | BLA_CAT_G137_PDZ | Chloramphenicol acetyltransferase with a PDZ insertion behind G137 |
| 12 | BLA_CAT_W150_PDZ | Chloramphenicol acetyltransferase with a PDZ insertion behind W150 |
| 13 | BLA_CAT_V160_PDZ | Chloramphenicol acetyltransferase with a PDZ insertion behind V160 |
| 14 | BLA_CAT_A161_PDZ | Chloramphenicol acetyltransferase with a PDZ insertion behind A161 |
| 15 | BLA_CAT_N162_PDZ | Chloramphenicol acetyltransferase with a PDZ insertion behind N162 |
| 16 | BLA_CAT_D181_PDZ | Chloramphenicol acetyltransferase with a PDZ insertion behind D181 |
| 17 | BLA_CAT_Q190_PDZ | Chloramphenicol acetyltransferase with a PDZ insertion behind Q190 |
| 18 | BLA_CAT_F25_LOV | Chloramphenicol acetyltransferase with a AsLOV2 insertion behind F25 |
| 19 | BLA_CAT_Q26_LOV | Chloramphenicol acetyltransferase with a AsLOV2 insertion behind Q26 |
| 20 | BLA_CAT_S27_LOV | Chloramphenicol acetyltransferase with a AsLOV2 insertion behind S27 |
| 21 | BLA_CAT_T36_LOV | Chloramphenicol acetyltransferase with a AsLOV2 insertion behind T36 |
| 22 | BLA_CAT_F73_LOV | Chloramphenicol acetyltransferase with a AsLOV2 insertion behind F73 |
| 23 | BLA_CAT_C91_LOV | Chloramphenicol acetyltransferase with a AsLOV2 insertion behind C91 |
| 24 | BLA_CAT_P135_LOV | Chloramphenicol acetyltransferase with a AsLOV2 insertion behind P135 |
| 25 | BLA_CAT_K136_LOV | Chloramphenicol acetyltransferase with a AsLOV2 insertion behind K136 |
| 26 | BLA_CAT_G137_LOV | Chloramphenicol acetyltransferase with a AsLOV2 insertion behind G137 |
| 27 | BLA_CAT_W150_LOV | Chloramphenicol acetyltransferase with a AsLOV2 insertion behind W150 |
| 28 | BLA_CAT_V160_LOV | Chloramphenicol acetyltransferase with a AsLOV2 insertion behind V160 |
| 29 | BLA_CAT_A161_LOV | Chloramphenicol acetyltransferase with a AsLOV2 insertion behind A161 |
| 30 | BLA_CAT_N162_LOV | Chloramphenicol acetyltransferase with a AsLOV2 insertion behind N162 |
| 31 | BLA_CAT_D181_LOV | Chloramphenicol acetyltransferase with a AsLOV2 insertion behind D181 |
| 32 | BLA_CAT_Q190_LOV | Chloramphenicol acetyltransferase with a AsLOV2 insertion behind Q190 |
| 33 | BLA_CAT_K136_GPG_LOV2 | Chloramphenicol acetyltransferase with a AsLOV2 insertion behind K136, flanked by GPG linkers |
| 34 | eGFP | CMV promoter, eGFP, bGH-polyA |
| 35 | PAC_eGFP | CMV promoter, puromycin acetyltransferase, T2A sequence, eGFP, bGH-polyA |
| 36 | PAC_D29_PDZ_eGFP | CMV promoter, puromycin acetyltransferase with a PDZ insertion behind D29, T2A sequence, eGFP, bGH-polyA |
| 37 | PAC_G58_PDZ_eGFP | CMV promoter, puromycin acetyltransferase with a PDZ insertion behind G58, T2A sequence, eGFP, bGH-polyA |
| 38 | PAC_V75_PDZ_eGFP | CMV promoter, puromycin acetyltransferase with a PDZ insertion behind V75, T2A sequence, eGFP, bGH-polyA |
| 39 | PAC_S81_PDZ_eGFP | CMV promoter, puromycin acetyltransferase with a PDZ insertion behind S81, T2A sequence, eGFP, bGH-polyA |
| 40 | PAC_V82_PDZ_eGFP | CMV promoter, puromycin acetyltransferase with a PDZ insertion behind V82, T2A sequence, eGFP, bGH-polyA |
| 41 | PAC_E83_PDZ_eGFP | CMV promoter, puromycin acetyltransferase with a PDZ insertion behind E83, T2A sequence, eGFP, bGH-polyA |
| 42 | PAC_A84_PDZ_eGFP | CMV promoter, puromycin acetyltransferase with a PDZ insertion behind A84, T2A sequence, eGFP, bGH-polyA |
| 43 | PAC_G85_PDZ_eGFP | CMV promoter, puromycin acetyltransferase with a PDZ insertion behind G85, T2A sequence, eGFP, bGH-polyA |
| 44 | PAC_A86_PDZ_eGFP | CMV promoter, puromycin acetyltransferase with a PDZ insertion behind A86, T2A sequence, eGFP, bGH-polyA |
| 45 | PAC_S101_PDZ_eGFP | CMV promoter, puromycin acetyltransferase with a PDZ insertion behind S101, T2A sequence, eGFP, bGH-polyA |
| 46 | PAC_G111_PDZ_eGFP | CMV promoter, puromycin acetyltransferase with a PDZ insertion behind G111, T2A sequence, eGFP, bGH-polyA |
| 47 | PAC_K119_PDZ_eGFP | CMV promoter, puromycin acetyltransferase with a PDZ insertion behind K119, T2A sequence, eGFP, bGH-polyA |
| 48 | PAC_E120_PDZ_eGFP | CMV promoter, puromycin acetyltransferase with a PDZ insertion behind E120, T2A sequence, eGFP, bGH-polyA |
| 49 | PAC_P121_PDZ_eGFP | CMV promoter, puromycin acetyltransferase with a PDZ insertion behind P121, T2A sequence, eGFP, bGH-polyA |

| | | |
|---|---|---|
| **50** | PAC_P185_PDZ_eGFP | CMV promoter, puromycin acetyltransferase with a PDZ insertion behind P185, T2A sequence, eGFP, bGH-polyA |
| **51** | PAC_E83_LOV_eGFP | CMV promoter, puromycin acetyltransferase with a AsLOV2 insertion behind E83, T2A sequence, eGFP, bGH-polyA |
| **52** | Renilla luciferase | HSV TK promoter, Renilla luciferase, SV40 polyA (Promega) |
| **53** | TetO_firefly_luciferase[6] | TetO, firefly luciferase |
| **54** | sgRNA-Tet_RLuc (SpCas9)[7] | U6 promoter, TetO-targeting sgRNA for MbCas12a, polyT; HSV TK promoter, Renilla luciferase, SV40 polyA |
| **55** | d*Sp*Cas9-VPR | EF-1α promoter, SV40 NLS, d*Sp*Cas9, SV40 NLS, VPR, beta-globin polyA |
| **56** | d*Sp*Cas9-N309-LOV-VPR | EF-1α promoter, SV40 NLS, d*Sp*Cas9 with an AsLOV2 insertion behind N309, SV40 NLS, VPR, beta-globin polyA |
| **57** | d*Sp*Cas9-E566-LOV-VPR | EF-1α promoter, SV40 NLS, d*Sp*Cas9 with an AsLOV2 insertion behind E566, SV40 NLS, VPR, beta-globin polyA |
| **58** | d*Sp*Cas9-T1048-LOV-VPR | EF-1α promoter, SV40 NLS, d*Sp*Cas9 with an AsLOV2 insertion behind T1048, SV40 NLS, VPR, beta-globin polyA |
| **59** | d*Sp*Cas9-D1284-LOV-VPR | EF-1α promoter, SV40 NLS, d*Sp*Cas9 with an AsLOV2 insertion behind D1284, SV40 NLS, VPR, beta-globin polyA |
| **60** | Firefly luciferase[8] | TK promoter, firefly luciferase |
| **61** | nt-sgRNA (*Mb*Cas12a) | U6 promoter, non-targeting sgRNA for MbCas12a, polyT |
| **62** | ff-sgRNA (*Mb*Cas12a) | U6 promoter, firefly luciferase-targeting sgRNA for MbCas12a, polyT |
| **63** | MbCas12a[9] | CMV promoter, MbCas12a, nucleoplasmin NLS, beta-globin polyA |
| **64** | MbCas12a_F35_PDZ | CMV promoter, MbCas12a with a PDZ insertion behind F35, nucleoplasmin NLS, beta-globin polyA |
| **65** | MbCas12a_L36_PDZ | CMV promoter, MbCas12a with a PDZ insertion behind L36, nucleoplasmin NLS, beta-globin polyA |
| **66** | MbCas12a_N37_PDZ | CMV promoter, MbCas12a with a PDZ insertion behind N37, nucleoplasmin NLS, beta-globin polyA |
| **67** | MbCas12a_T71_PDZ | CMV promoter, MbCas12a with a PDZ insertion behind T71, nucleoplasmin NLS, beta-globin polyA |
| **68** | MbCas12a_K72_PDZ | CMV promoter, MbCas12a with a PDZ insertion behind K72, nucleoplasmin NLS, beta-globin polyA |
| **69** | MbCas12a_L73_PDZ | CMV promoter, MbCas12a with a PDZ insertion behind L73, nucleoplasmin NLS, beta-globin polyA |
| **70** | MbCas12a_N113_PDZ | CMV promoter, MbCas12a with a PDZ insertion behind N113, nucleoplasmin NLS, beta-globin polyA |
| **71** | MbCas12a_G114_PDZ | CMV promoter, MbCas12a with a PDZ insertion behind G114, nucleoplasmin NLS, beta-globin polyA |
| **72** | MbCas12a_G115_PDZ | CMV promoter, MbCas12a with a PDZ insertion behind G115, nucleoplasmin NLS, beta-globin polyA |
| **73** | MbCas12a_I486_PDZ | CMV promoter, MbCas12a with a PDZ insertion behind I486, nucleoplasmin NLS, beta-globin polyA |
| **74** | MbCas12a_K487_PDZ | CMV promoter, MbCas12a with a PDZ insertion behind K487, nucleoplasmin NLS, beta-globin polyA |
| **75** | MbCas12a_S488_PDZ | CMV promoter, MbCas12a with a PDZ insertion behind S488, nucleoplasmin NLS, beta-globin polyA |
| **76** | MbCas12a_P1152_PDZ | CMV promoter, MbCas12a with a PDZ insertion behind P1152, nucleoplasmin NLS, beta-globin polyA |
| **77** | MbCas12a_N1153_PDZ | CMV promoter, MbCas12a with a PDZ insertion behind N1153, nucleoplasmin NLS, beta-globin polyA |
| **78** | MbCas12a_L1154_PDZ | CMV promoter, MbCas12a with a PDZ insertion behind L1154, nucleoplasmin NLS, beta-globin polyA |
| **79** | MbCas12a_D55_PDZ | CMV promoter, MbCas12a with a PDZ insertion behind D55, nucleoplasmin NLS, beta-globin polyA |
| **80** | MbCas12a_K105_PDZ | CMV promoter, MbCas12a with a PDZ insertion behind K105, nucleoplasmin NLS, beta-globin polyA |
| **81** | MbCas12a_I426_PDZ | CMV promoter, MbCas12a with a PDZ insertion behind I426, nucleoplasmin NLS, beta-globin polyA |
| **82** | MbCas12a_L501_PDZ | CMV promoter, MbCas12a with a PDZ insertion behind L501, nucleoplasmin NLS, beta-globin polyA |
| **83** | MbCas12a_A802_PDZ | CMV promoter, MbCas12a with a PDZ insertion behind A802, nucleoplasmin NLS, beta-globin polyA |
| **84** | MbCas12a_K487_GP_AsLOV2 | CMV promoter, MbCas12a with a AsLOV2 insertion behind K487 flanked by a GP linker, nucleoplasmin NLS, beta-globin polyA |
| **85** | MbCas12a_N1153_GP_AsLOV2 | CMV promoter, MbCas12a with a AsLOV2 insertion behind N1153 flanked by a GP linker, nucleoplasmin NLS, beta-globin polyA |
| **86** | MbCas12a_K487_GPG_GR2 | CMV promoter, MbCas12a with a cpGR2 insertion behind K487 flanked by a GPPG linker, nucleoplasmin NLS, beta-globin polyA |
| **87** | MbCas12a_N1153_GPG_GR2 | CMV promoter, MbCas12a with a cpGR2 insertion behind N1153 flanked by a GPPG linker, nucleoplasmin NLS, beta-globin polyA |
| **88** | VEGFA-sgRNA (*Mb*Cas12a) | U6 promoter, VEGFA targeting sgRNA for MbCas12a, polyT |
| **89** | RUNX-sgRNA (*Mb*Cas12a) | U6 promoter, RUNX targeting sgRNA for MbCas12a, polyT |
| **90** | GRIN2B-sgRNA (*Mb*Cas12a) | U6 promoter, GRIN2B targeting sgRNA for MbCas12a, polyT |

**Supplementary Table 2 | Amino acid sequences of the domains and proteins used in this study.** Blue: linker sequences; orange: affinity tag; red: T2A sequence; green: nuclear localization sequence.

| Protein / Domain | Amino acid sequence |
|---|---|
| Chloramphenicol acetyltransferase | MEKKITGYTTVDISQWHRKEHFEAFQSVAQCTYNQTVQLDITAFLKTVKKNKHKFY<br>PAFIHILARLMNAHPEFRMAMKDGELVIWDSVHPCYTVFHEQTETFSSLLSEYHDD<br>FRQFLHIYSQDVACYGENLAYFPKGFIENMFFVSANPWVSFTSFDLNVANMDNFF<br>APVFTMGKYYTQGDKVLMPLAIQVHHAVCDGFHVGRMLNELQQYCDEWQGGA |
| *As*LOV2 | SGLATTLERIEKNFVITDPRLPDNPIIFASDSFLQLTEYSREEILGRNCRFLQGPETD<br>RATVRKIRDAIDNQTEVTVQLINYTKSGKKFWNLFHLQPMRDQKGDVQYFIGVQLD<br>GTEHVRDAAEREGVMLIKKTAENIDEAAKGS |
| PDZ | SGRRRVTVRKADAGGLGISIKGGRENKMPILISKIFKGLAADQTEALFVGDAILSVN<br>GEDLSSATHDEAVQALKKTGKEVVLEVKYMKGS |
| GR2 | GPGNSSQNWQRFYQLTKLLDSMHEMVGGLLQFCFYTFVNKSLSVEFPEMLAEIIS<br>NQLPKFKAGSVKPLLFHQKGGGSGGSGGSGGSGGSGLISLLEVIEPEVLYSGYDS<br>TLPDTSTRLMSTLNRLGGRQVVSAVKWAKALPGFRNLHLDDQMTLLQYSWMSLM<br>AFSLGWRSYKQSNGNMLCFAPDLVINEERMQLPYMYDQCQQMLKISSEFVRLQV<br>SYDEYLCMKVLLLLSTVPKDGLKSQAVFDEIRMTYIKELGKAIVKREGGPG |
| Puromycin acetyltransferase-T2A-eGFP | MTEYKPTVRLATRDDVPRAVRTLAAAFADYPATRHTVDPDRHIERVTELQELFLTR<br>VGLDIGKVWVADDGAAVAVWTTPESVEAGAVFAEIGPRMAELSGSRLAAQQQME<br>GLLAPHRPKEPAWFLATVGVSPDHQGKGLGSAVVLPGVEAAERAGVPAFLETSA<br>PRNLPFYERLGFTVTADVEVPEGPRTWCMTRKPGAGSTGSRGSGEGRGSLLTC<br>GDVEENPGPMVSKGEELFTGVVPILVELDGDVNGHKFSVSGEGEGDATYGKLTLK<br>FICTTGKLPVPWPTLVTTLTYGVQCFSRYPDHMKQHDFFKSAMPEGYVQERTIFF<br>KDDGNYKTRAEVKFEGDTLVNRIELKGIDFKEDGNILGHKLEYNYNSHNVYIMADK<br>QKNGIKVNFKIRHNIEDGSVQLADHYQQNTPIGDGPVLLPDNHYLSTQSALSKDPN<br>EKRDHMVLLEFVTAAGITLGMDELYK |
| *Mb*Cas12a | MLFQDFTHLYPLSKTVRFELKPIGKTLEHIHAKNFLNQDETMADMYQKVKAILDDY<br>HRDFIADMMGEVKLTKLAEFYDVYLKFRKNPKDDGLQKQLKDLQAVLRKEIVKPIG<br>NGGKYKAGYDRLFGAKLFKDGKELGDLAKFVIAQEGESSPKLAHLAHFEKFSTYFT<br>GFHDNRKNMYSDEDKHTAIAYRLIHENLPRFIDNLQILATIKQKHSALYDQIINELTA<br>SGLDVSLASHLDGYHKLLTQEGITAYNTLLGGISGEAGSRKIQGINELINSHHNQHC<br>HKSERIAKLRPLHKQILSDGMGVSFLPSKFADDSEVCQAVNEFYRHYADVFAKVQ<br>SLFDGFDDYQKDGIYVEYKNLNELSKQAFGDFALLGRVLDGYYVDVVNPEFNERF<br>AKAKTDNAKAKLTKEKDKFIKGVHSLASLEQAIEHYTARHDDESVQAGKLGQYFKH<br>GLAGVDNPIQKIHNNHSTIKGFLERERPAGERALPKIKSDKSPEIRQLKELLDNALN<br>VAHFAKLLTTKTTLHNQDGNFYGEFGALYDELAKIATLYNKVRDYLSQKPFSTEKY<br>KLNFGNPTLLNGWDLNKEKDNFGVILQKDGCYYLALLDKAHKKVFDNAPNTGKSV<br>YQKMIYKLLPGPNKMLPKVFFAKSNLDYYNPSAELLDKYAQGTHKKGDNFNLKDC<br>HALIDFFKAGINKHPEWQHFGFKFSPTSSYQDLSDFYREVEPQGYQVKFVDINADY<br>INELVEQGQLYLFQIYNKDFSPKAHGKPNLHTLYFKALFSEDNLVNPIYKLNGEAEIF<br>YRKASLDMNETTIHRAGEVLENKNPDNPKKRQFVYDIIKDKRYTQDKFMLHVPITM<br>NFGVQGMTIKEFNKKVNQSIQQYDEVNVIGIDRGERHLLYLTVINSKGEILEQRSLN<br>DITTASANGTQMTTPYHKILDKREIERLNARVGWGEIETIKELKSGYLSHVVHQISQ<br>LMLKYNAIVVLEDLNFGFKRGRFKVEKQIYQNFENALIKKLNHLVLKDKADDEIGSY<br>KNALQLTNNFTDLKSIGKQTGFLFYVPAWNTSKIDPETGFVDLLKPRYENIAQSQAF<br>FGKFDKICYNADRGYFEFHIDYAKFNDKAKNSRQIWKICSHGDKRYVYDKTANQN<br>KGATIGVNVNDELKSLFTRYHINDKQPNLVMDICQNNDKEFHKSLMYLLKTLLALRY<br>SNASSDEDFILSPVANDEGVFFNSALADDTQPQNADANGAYHIALKGLWLLNELKN<br>SDDLNKVKLAIDNQTWLNFAQNRKRPAATKKAGQAKKKKGSYPYDVPDYAYPYD<br>VPDYAYPYDVPDYA |
| d*Sp*Cas9-VPR | MSPKKKRKVEASDKKYSIGLAIGTNSVGWAVITDEYKVPSKKFKVLGNTDRHSIKK<br>NLIGALLFDSGETAEATRLKRTARRRYTRRKNRICYLQEIFSNEMAKVDDSFFHRLE<br>ESFLVEEDKKHERHPIFGNIVDEVAYHEKYPTIYHLRKKLVDSTDKADLRLIYLALAH<br>MIKFRGHFLIEGDLNPDNSDVDKLFIQLVQTYNQLFEENPINASGVDAKAILSARLS<br>KSRRLENLIAQLPGEKKNGLFGNLIALSLGLTPNFKSNFDLAEDAKLQLSKDTYDDD<br>LDNLLAQIGDQYADLFLAAKNLSDAILLSDILRVNTEITKAPLSASMIKRYDEHHQDL<br>TLLKALVRQQLPEKYKEIFFDQSKNGYAGYIDGGASQEEFYKFIKPILEKMDGTEEL<br>LVKLNREDLLRKQRTFDNGSIPHQIHLGELHAILRRQEDFYPFLKDNREKIEKILTFRI<br>PYYVGPLARGNSRFAWMTRKSEETITPWNFEEVVDKGASAQSFIERMTNFDKNLP<br>NEKVLPKHSLLYEYFTVYNELTKVKYVTEGMRKPAFLSGEQKKAIVDLLFKTNRKV<br>TVKQLKEDYFKKIECFDSVEISGVEDRFNASLGTYHDLLKIIKDKDFLDNEENEDILE<br>DIVLTLTLFEDREMIEERLKTYAHLFDDKVMKQLKRRRYTGWGRLSRKLINGIRDK<br>QSGKTILDFLKSDGFANRNFMQLIHDDSLTFKEDIQKAQVSGQGDSLHEHIANLAG |

| | |
|---|---|
| | SPAIKKGILQTVKVVDELVKVMGRHKPENIVIEMARENQTTQKGQKNSRERMKRIE<br>EGIKELGSQILKEHPVENTQLQNEKLYLYYLQNGRDMYVDQELDINRLSDYDVDAI<br>VPQSFLKDDSIDNKVLTRSDKNRGKSDNVPSEEVVKKMKNYWRQLLNAKLITQRK<br>FDNLTKAERGGLSELDKAGFIKRQLVETRQITKHVAQILDSRMNTKYDENDKLIREV<br>KVITLKSKLVSDFRKDFQFYKVREINNYHHAHDAYLNAVVGTALIKKYPKLESEFVY<br>GDYKVYDVRKMIAKSEQEIGKATAKYFFYSNIMNFFKTEITLANGEIRKRPLIETNGE<br>TGEIVWDKGRDFATVRKVLSMPQVNIVKKTEVQTGGFSKESILPKRNSDKLIARKK<br>DWDPKKYGGFDSPTVAYSVLVVAKVEKGKSKKLKSVKELLGITIMERSSFEKNPID<br>FLEAKGYKEVKKDLIIKLPKYSLFELENGRKRMLASAGELQKGNELALPSKYVNFLY<br>LASHYEKLKGSPEDNEQKQLFVEQHKHYLDEIIEQISEFSKRVILADANLDKVLSAY<br>NKHRDKPIREQAENIIHLFTLTNLGAPAAFKYFDTTIDRKRYTSTKEVLDATLIHQSIT<br>GLYETRIDLSQLGGDSAGGGGSGGGGSGGGGSGPKKKRKVAAAGSGRADALDD<br>FDLDMLGSDALDDFDLDMLGSDALDDFDLDMLGSDALDDFDLDMLINTSGGSGG<br>GSGGSSQYLPDTDDRHRIEEKRKRTYETFKSIMKKSPFSGPTDPRPPPRRIAVPSR<br>SSASVPKPAPQPYPFTSSLSTINYDEFPTMVFPSGQISQASALAPAPPQVLPQAPA<br>PAPAPAMVSALAQAPAPVPVLAPGPPQAVAPPAPKPTQAGEGTLSEALLQLQFDD<br>EDLGALLGNSTDPAVFTDLASVDNSEFQQLLNQGIPVAPHTTEPMLMEYPEAITRL<br>VTGAQRPPDPAPAPLGAPGLPNGLLSGDEDFSSIADMDFSALLGSGSGSRDSRE<br>GMFLPKPEAGSAISDVFEGREVCQPKRIRPFHPPGSPWANRPLPASLAPTPTGPV<br>HEPVGSLTPAPVPQPLDPAPAVTPEASHLLEDPDEDTSQAVKALREMADTVIPQK<br>EEAAICGQMDLSHPPPRGHLDELTTTLESMTEDLNLDSPLTPELNEILDTFLNDECL<br>LHAMHISTGLSIFDTSLFG |

**Supplementary Table 3 | gRNAs used in this study.** Spacer sequences are marked in bold, PAM motifs are underlined.

| Target | Sequence 5'-3' | Cas protein | Source |
|---|---|---|---|
| *RUNX* | TTTA**CCTTCGGAGCGAAAACCAAG** | *Mb*Cas12a | [9] |
| *GRIN2B* | TTTG**GTGCTCAATGAAAGGAGATAAGG** | *Mb*Cas12a | [9] |
| *VEGFA* | TTTG**CTAGGAATATTGAAGGGGGC** | *Mb*Cas12a | [10] |
| Firefly luciferase | TTTC**GGCAACCAGATCATCCCCGACAC** | *Mb*Cas12a | This work |
| TetO | **TCTCTATCACTGATAGGGAG**TGG | *Sp*Cas9 | [7] |

**Supplementary Table 4 | Primers for amplification of genomic loci.**

| Gene | Orientation | Primer sequence 5'-3' |
|---|---|---|
| *RUNX* | Forward | CCAGAGGTATCCAGCAGAGG |
| | Reverse | TACAGGCAAAGCTGAGCAAA |
| *GRIN2B* | Forward | GTGTATGCATACTCGCATGGC |
| | Reverse | CAGGACGGCCAACACCAAC |
| *VEGFA* | Forward | GGGTCACTCCAGGATTCCAATAG |
| | Reverse | GCAATGAAGGGGAAGCTCGAC |
